# Lifespan Trajectories of Resting State EEG power

**DOI:** 10.64898/2026.06.02.729638

**Authors:** David B. Chorlian, Jacquelyn L. Meyers, Andrey Anokhin, Chella Kamarajan, Ashwini K. Pandey, Jian Zhang, Sivan Kinreich, Bernice Porjesz

## Abstract

Trajectories of resting state monopolar EEG power were analyzed using the means, variances, and correlations of data in seven frequency bands obtained from 17 electrodes of the 10-20 system from a sample of 5238 participants from the Collaborative Study on the Genetics of Alcoholism (COGA) with 11906 observations from ages 12 through 70. The values for the individual observations were calculated by standard Fourier transform based methods. In order to make the study more useful for understanding EEG in the general population only subjects who were never diagnosed with alcohol use disorder were included in the study. The trajectories of power show a clear pattern of decrease in all frequency bands and both sexes from ages 12 to 20. Subsequently there is considerable amount of variation between frequency bands and between males and females. In females, the generally lower rate of decrease after age 20 is reversed at age 35 in anterior and central regions in the alpha and beta bands, and stabilized in the theta bands. In males, the decreases continue in the theta and high alpha bands, but there are elements of the reversal in the low alpha and beta bands. Examination of the derivatives of the trajectories show that sex differences begin in the mid-twenties in alpha and beta but not until the mid-thirties in theta.

In contrast to the varied power value trajectories, the trajectories of inter-frequency correlation show a pervasive increase with age of the correlation between high alpha and each of the other frequency bands except high beta. This increase is primarily anterior in the alpha-theta correlations and more regionally uniform in the alpha-beta correlations. Sex differences are very small. Trajectories of intra-frequency correlations for comparable between region pairs and within region pairs were generally high and stable across age. This is the first of a series of studies which will provide similar analyses of bipolar EEG power and bipolar EEG coherence in this sample. We know of no other study of resting state EEG power which combines the analysis of power with the analysis of both inter-frequency and intra-frequency correlation of power values.

## 1 Introduction

The Collaborative Study on the Genetics of Alcoholism (COGA) (Begleiter et al., 1995; Agrawal et al., 2023; Meyers et al., 2023) has published many studies of resting state EEG since its inception covering both power and coherence primarily related to alcohol related conditions (Rangaswamy et al., 2002, 2003), but also genetic aspects (Porjesz et al., 2002; Tang et al., 2007a,b; Chorlian et al., 2007), and combinations of these factors (Meyers et al., 2017, 2019, 2021a,b; Chorlian et al., 2024). COGA is a participant in the ENIGMA-EEG working group and involved in projects described in Smit et al. (2018) and Smit et al. (2021). Continuing in the theme of those projects we performed a comprehensive analysis of resting state EEG power values extending from age 12 to age 70, which included estimations of mean values and of both inter-frequency and intra-frequency correlations, using methods similar to those used in our previous studies as noted above, and described in detail in Chorlian et al. (2015) and Chorlian et al. (2024). Specifically, standard Fourier transform methods were used to estimate power in 1/2 Hz bins and aggregated in seven frequency bands used in previous studies. This data was then subject to a non-parametric regression analysis and complementary covariance estimation at 117 1/2 year age centers from age 12 to age 70.

We observed a variety of features, primarily in our correlation results, which we believe are previously unreported in the EEG literature. We note that our observations are not uniformly distributed across the analyzed age range, being highly skewed toward the younger ages, we thought it desirable to present these because the effective sample sizes were large compared to many EEG studies, as explained in section 2.3.1. We used conventional frequency band based analysis in order to make possible comparisons to earlier studies. This is the first of series of studies using an identical methodology; subsequent studies will cover bipolar EEG power and bipolar EEG coherence.

## 2 Materials and Methods

### 2.1 Sample

As part of the Collaborative Study on the Genetics of Alcoholism (COGA) (Begleiter et al., 1995; **?**), EEG data were collected from families densely affected with alcohol use disorder (AUD) and their offspring as well as community comparison families. Many of the initially recruited subjects were recontacted recently and additional EEG data was collected. This provides a sample of subjects with observations ranging from age 10 to over 70 years of age. A phase of the study in which the offspring of previous participants were tested repeatedly from ages 12 and onward at approximately 2 year intervals provided a large number of observations from individuals under age 30. In order to make this study more comparable to results from the general population, all data from subjects who at any age had a DSM-5 AUD diagnosis were removed from the study. Subjects were not excluded for other alcohol use behavior.

### 2.2 Data Collection and Processing

Data was collected as described in Chorlian et al. (2009) and Meyers et al. (2019). Subjects were asked to keep their eyes closed and not to move their head while data from 17 electrodes of the 10-20 system was recorded for a period of 256 seconds in an enclosed booth. Data cleaning was effected by a subspace removal method, removing a portion of the data from the scalp channels correlated to the eye channel. The one second segments in which the maximal difference among observations at any one electrode exceeded 100 micro volts were marked as unusable. Subject data without at least 129 usable segments was rejected.

Power was calculated using a variation of the method used in Smit et al. (2018). Cleaned data were imported into MATLAB, organized into overlapping 2 s epochs, detrended, and power spectra calculated using Fast Fourier Transformation (FFT). Power was defined as the squared radius of the orthogonal sine and cosine amplitudes averaged over window size, and the mean value taken for the frequency band to obtain power density. Power values were aggregated into seven frequency bands from low theta to high beta and log transformed to obtain approximate normality. Details of the frequency bands and the electrodes used may be found in section 4.

### 2.3 Analysis Methods

#### 2.3.1 Regression Modeling

A non-parametric local linear regression (LOESS) method was used to estimate the mean and variance of the power data at 117 age centers 1/2 year apart for ages from 12 to 70. The bandwidth for the kernel (Epanechnikov) was taken as 0.6. More specifically, the model for each age *t*_*k*_, *k* = 1 … 117, is represented by the following equation, in which *y*_*i j*_ is the dependent variable for subject *i* at observation *j* at age *t*_*i j*_, *W*_*h*_(*t*_*i j*_ − *t*_*k*_) the kernel weighting for the local linear regression model, a function of the bandwidth (fraction of data included) *h, β*_0,*n,k*_ the parameter to be estimated for covariate *n* for its effect on the intercept and *β*_1,*n,k*_ the parameter to be estimated for covariate *n* for its effect on the slope, and *X*_*i, j,m*_ the value of covariate *m* of subject *i* at observation *j*. Different sets of covariates may be used for the estimation of the slope and the intercept. The weight *W*_*h*_(*t*_*i j*_− *t*_*k*_) will be non-zero only for a set of values near *t*_*k*_. Note that the beta values are indexed to emphasize that each is the product of a separate estimation for each age. (In the equation, intercept terms appear on the first line and slope terms on the second line. The summations are over the repeated indices in the covariates and betas.)

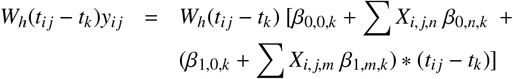

In the statistical model gender is the only covariate for the intercept and slope. Weights for individuals were adjusted to account for multiple observations on single individuals. Results are not independent between models for different ages since the data in each of the 117 regression models has considerable overlap with the data used in the models for nearby ages. This model was first fully described in Chorlian et al. (2015). Unlike in its previous uses, the age distribution of the sample is skewed, with about 3/4 of the observations from subjects thirty years of age or younger. Although 60%of the observations are used in each regression calculation, the values of the weights depends on the distribution of the differences in age from the age center of the observations, and thus the skewed age distribution affects the age distribution of the effective sample size, defined as (∑*W*_*h*_(*t*_*i j*_ − *t*_*k*_))^2^*/* ∑(*W*_*h*_(*t*_*i j*_ −*t*_*k*_))^2^ for each *t*_*k*_, as illustrated in figure 11.

To fully characterize the sex differences in the power trajectories, derivatives of the trajectories were calculated by fitting a spline to the trajectory values, differentiating the spline polynomials, and smoothing the resulting values.

#### 2.3.2 Covariance Modeling

An adaptation of the regression method, initially described in Chorlian et al. (2015), was used calculate covariances. Briefly, the weights used in the regression calculation were employed in a weighted covariance analysis. The covariance of the data both between all pairs of electrodes for each frequency band and between all pairs of frequencies for each electrode was calculated for each of the age centers. In the covariance analysis males and females were analyzed separately, in contrast to the mean and variance analysis which uses covariates to estimate male and female trajectories. The use of covariates to estimate sex effects provides more accurate estimates of sex differences than separate analyses.

Correlational estimates derived from the covariance matrices rather than the covariances matrices themselves were analyzed in order to more clearly differentiate changes in power relations from changes in power values. With regard to the intra-frequency correlations, the electrodes fell into four groups by scalp position: frontal, central, parietal, and occipital, with 5 electrodes in each of the first three groups and 2 in the final group. In some cases the parietal and occipital electrodes were combined into the posterior group, in which case the frontal group is renamed the anterior group. This geometry means that the occipital electrodes were omitted from some correlational analyses because there could be no direct comparison with the other groups. Although some left/right differences were observed, there is no formal left/right analysis presented because of the greater variation along the anterior/posterior axis, with a secondary variation between peripheral electrodes and interior electrodes being clearly visible.

## 3 Results

Results are derived from visual inspection of the figures found in the plots found in section 5 and interpreted with consideration of the coefficients of variation obtained from the variance estimates of the analysis. The description is based on both the trajectories themselves and their rates of change (time derivatives) and on the trajectories of the inter-frequency electrode specific correlations. In many cases, changes in the shapes of the power trajectories are more easily identified by the changes in derivatives. The trajectories represent the cumulative effects of the entire developmental/aging processes; the derivatives track the age specific variations in the processes. When the derivatives are greater than zero, the trajectories are increasing, when less than zero, the trajectories are decreasing, The zero crossings of the derivative trajectories indicate a transition between increasing and decreasing processes. Extrema of the derivatives mark the greatest intensities of the processes.

The major results of the study are that there is large age variation in the power values between the theta bands and the alpha bands during the initial developmental stage, ages 12 to 20, with strikingly different patterns in both power and inter-frequency correlation, which continue into maturity. From the start (age 12) there are sex differences in the trajectories of high alpha. By age 25 sex differences extend to low alpha and beta bands but not until 35 in the theta bands. Then follows a maturation phase extending to age 50, and then a significant change into the aging process from age 50 to age 70 which has fewer short term fluctuations. The relative stability of the period from age 50 to age 70 is undoubtedly affected by the relative sparsity of observations (fewer than 1000) from subjects at ages greater than 50.

Inter-frequency correlation is characterized by the pervasive increase with age of the correlation of high alpha power with the power of all the other frequency bands except for high beta. In contrast, intra-frequency correlations are relatively stable over the entire age range and are similar in the different frequency bands. This suggests that volume conduction effects produce much of the observed correlation, also exhibited in the intra-modal theta band correlations illustrated in figure 2 of Chorlian et al. (2015).

Examination of the data suggests that there are initially distinct theta and alpha producing systems, with the theta system relatively homogeneous across brain regions from the frontal to the occipital and the alpha system with systematic differences between these brain regions. These observations are consistent with earlier well known work showing spatial variability of sources for activity in the alpha bands and the exception of alpha from the 1/f power law evidenced in the theta and beta band trajectories (Buszaki, 2006; Gasser et al., 1988; Röschke et al., 1997),

### 3.1 Trajectories of lifetime EEG power

Power trajectories are shown in figures 1 and 2. Development and aging over the course of the lifetime is characterized by considerable decreases in power from age 12 to age 20 in all frequency bands. In this age range the developmental pattern of trajectories of theta and alpha bands is quite dissimilar, with faster decreases in theta bands than in alpha bands, as well as frequency specific regional differences in the distribution of power values. The variation in power across scalp locations was very small in low theta and very large in high alpha. The variation in power in the theta band was primarily between central scalp locations and peripheral locations, while the variation in the alpha band was primarily between anterior and posterior locations. By the age of 30, all low theta values were less than 90% of their values at age 12, while no high alpha values were less than 90% of their values at age 12; the majority of the central and posterior electrodes had values greater than 95% of their initial values. In this period female values are generally larger than male values, and the rate of change (decrease) is larger in males than in females. Subsequently there are a variety of frequency and region dependent trajectories ranging from continued decreases in power in the theta bands to relative stability to increases, particularly in low alpha and in the beta frequency bands. Male power values were consistently lower than female values, but the shapes of the trajectories were similar from ages 12 to the early twenties, except in the high alpha band. Coefficients of variation were considerably smaller for theta values (about .07) than for alpha (about .1) or beta (about .14). However, in comparing values within frequency bands, the effective coefficients of variation should be reduced in value because of the relatively high levels of intra-frequency correlation.

**Figure 1:**
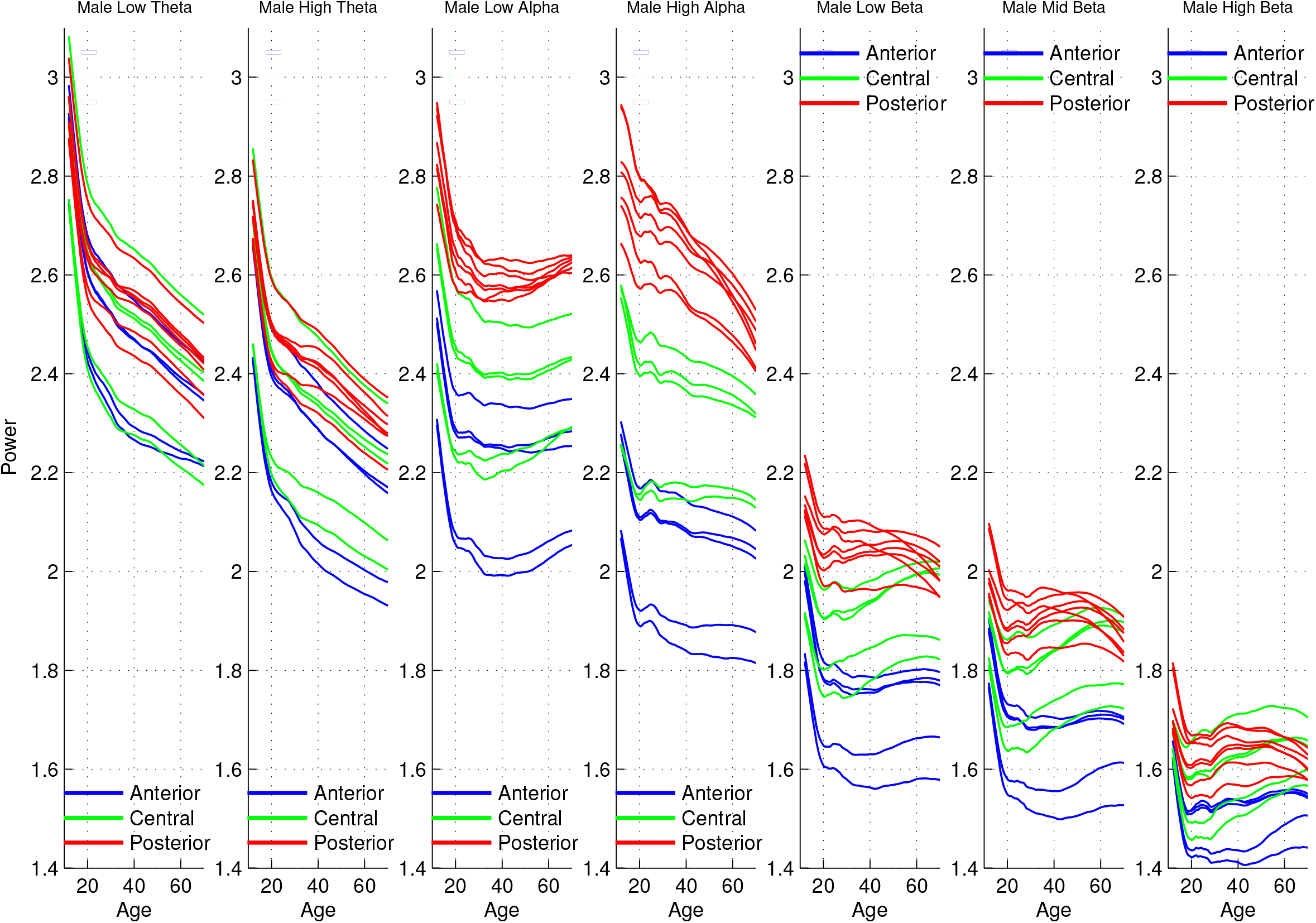
Trajectories of the seven frequency band male means color-coded by region.

**Figure 2:**
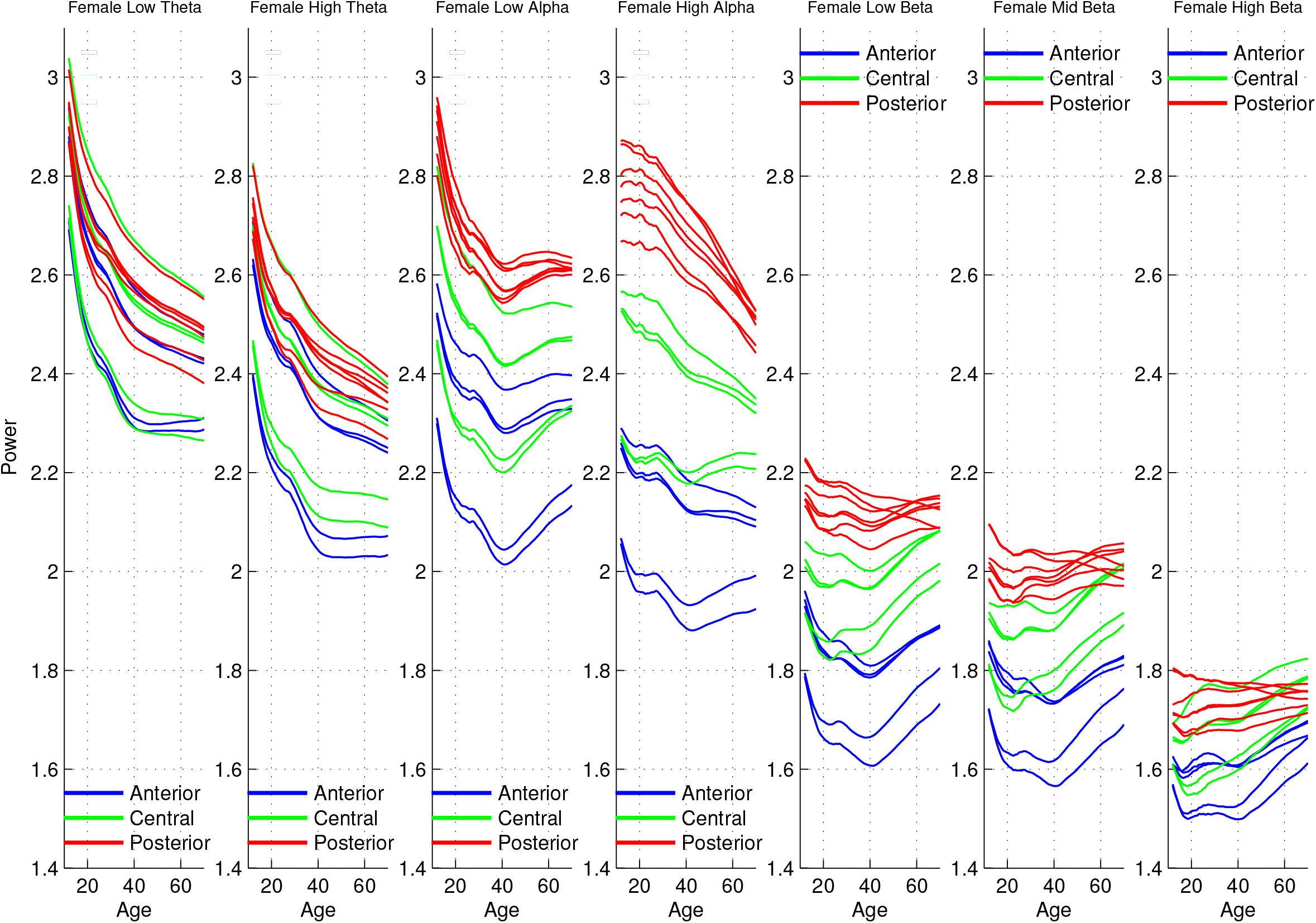
Trajectories of the seven frequency band female means color-coded by region.

The sex differences are much more apparent in the derivative plots. Figure 3 shows the lifetime derivative trajectories for the entire lifespan for both males and females. To age 20, there is little sex difference in the shapes of the derivatives. From age 20 to about 30 there are adjustments to the rate of decrease identifiable by local maxima in the derivatives, representing some adjustment after the large decreases in value before age 20. For females, after age 30 in every frequency band there is almost always a single succeeding local maximum for each electrode before age 50. For males, in every frequency band there are almost always two succeeding local maxima for each electrode before age 50. There variation in frequency specific and regional specific characteristics of these patterns as can be observed in figures 4 and figure 5.

**Figure 3:**
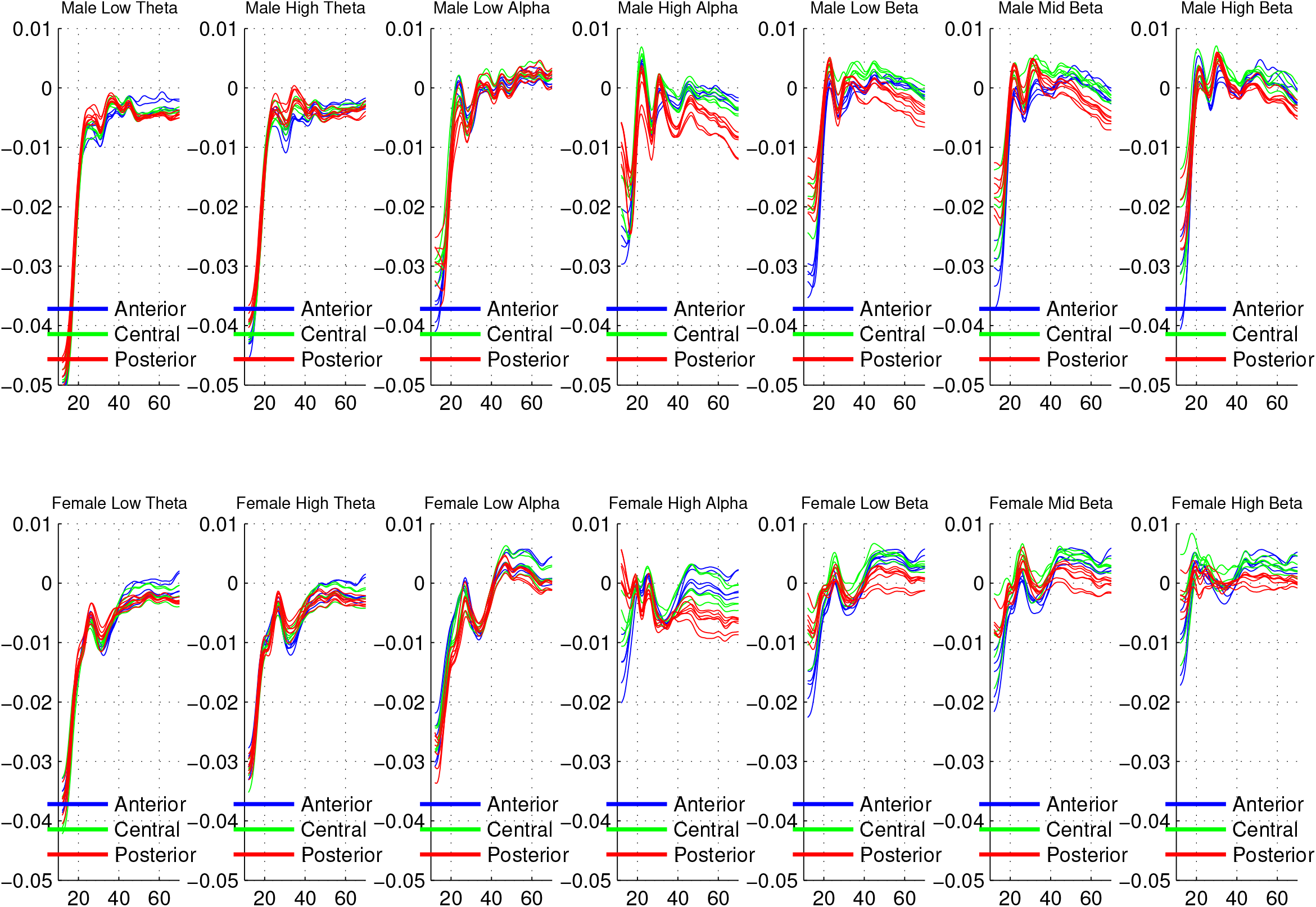
Derivatives of the entire male and female lifespan trajectories.

**Figure 4:**
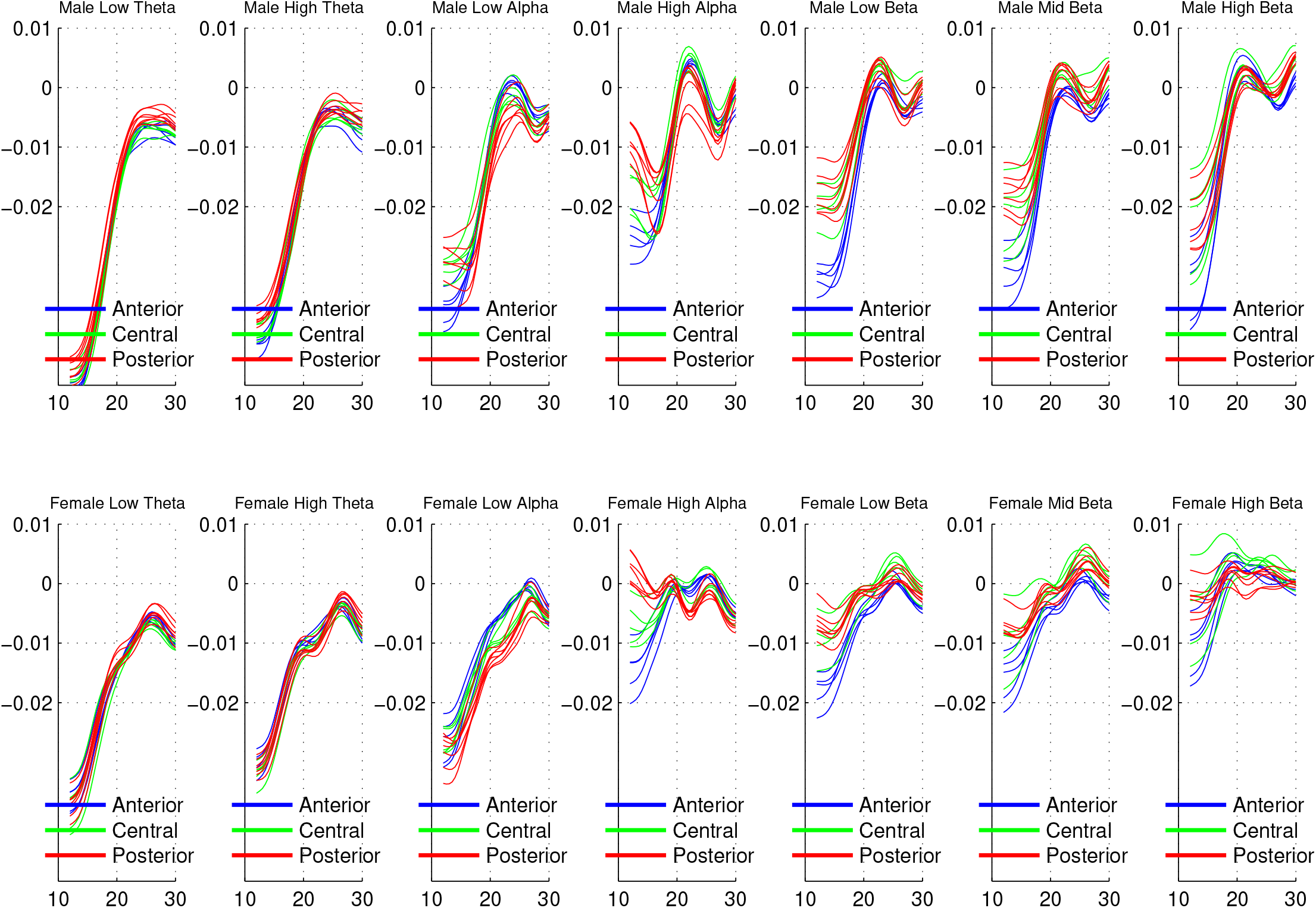
Derivatives of the male and female trajectories between age 12 and age 30. This is meant to contrast the general similarity between male and females with clear difference in the subsequent plot.

**Figure 5:**
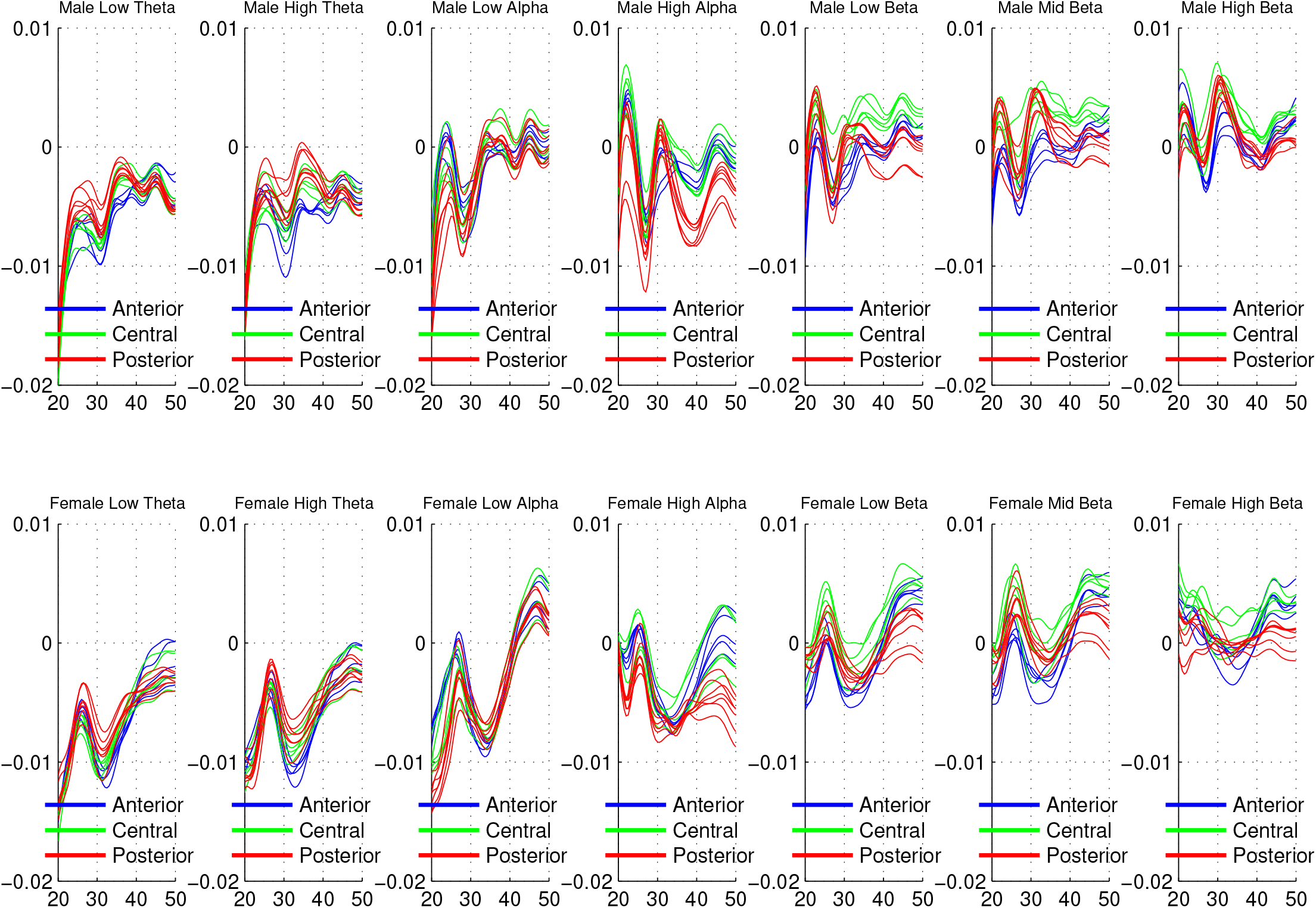
Derivatives of the entire male and female trajectories between ages 20 and 50. This is meant to contrast the general dissimilarity between male and females with general similarity in the previous plot.

### 3.2 Trajectories of lifetime EEG correlation

The age structures of inter-frequency correlations and intra-frequency correlations are quite different. While intra-frequency correlation trajectories are relatively stable over age and not different between frequency bands, inter-frequency correlations have a variety of trajectories which exhibit frequency and regional specificity. The most prominent feature is the pervasive increase with age of the correlation of high alpha power with the power of all the other frequency bands except for high beta. This increase is regionally specific for the alpha-theta correlations but not for the alpha-beta correlations. Most important in both the inter- and intra-frequency correlation trajectories is that there are no sharp decorrelations, and that relative values are generally preserved with age. This contrasts with the decorrelations observed in Chorlian et al. (2015) and non-linearity of the development of resting state EEG coherence suggested by the analysis performed in Chorlian et al. (2024). Additionally, divergence of trajectories is not a sign of decrease in correlation. This is most striking in the posterior alpha bands beginning at age 40 where the high alpha and low alpha trajectories have opposite slopes but increasing correlation.

#### 3.2.1 Inter-Frequency correlation

##### Theta-Alpha correlations

Inter-frequency correlations for the theta and alpha bands are shown in Figure 6. For this set of correlations in the theta and alpha frequency bands, arranged symmetrically with respect to the low theta and high alpha bands in this figure, there is considerable asymmetry between the low theta trajectories and the high alpha trajectories. The three rightmost correlations, all of which involve the high alpha band, show remarkable increases with age beginning from ages 20 to 30. The high alpha correlations also show considerable regional differences, not present to the same degree in the low theta correlations. In order to facilitate comparison with the alpha-beta correlations, in figure 7 the theta-alpha correlations are arranged in a row of 6 plots in which the three leftmost plots are adjacent frequency bands, the next two are those with one band separating the correlations, and finally the pair which is furthest apart. This figure shows clearly the strong regional differences in the high alpha correlations.

**Figure 6:**
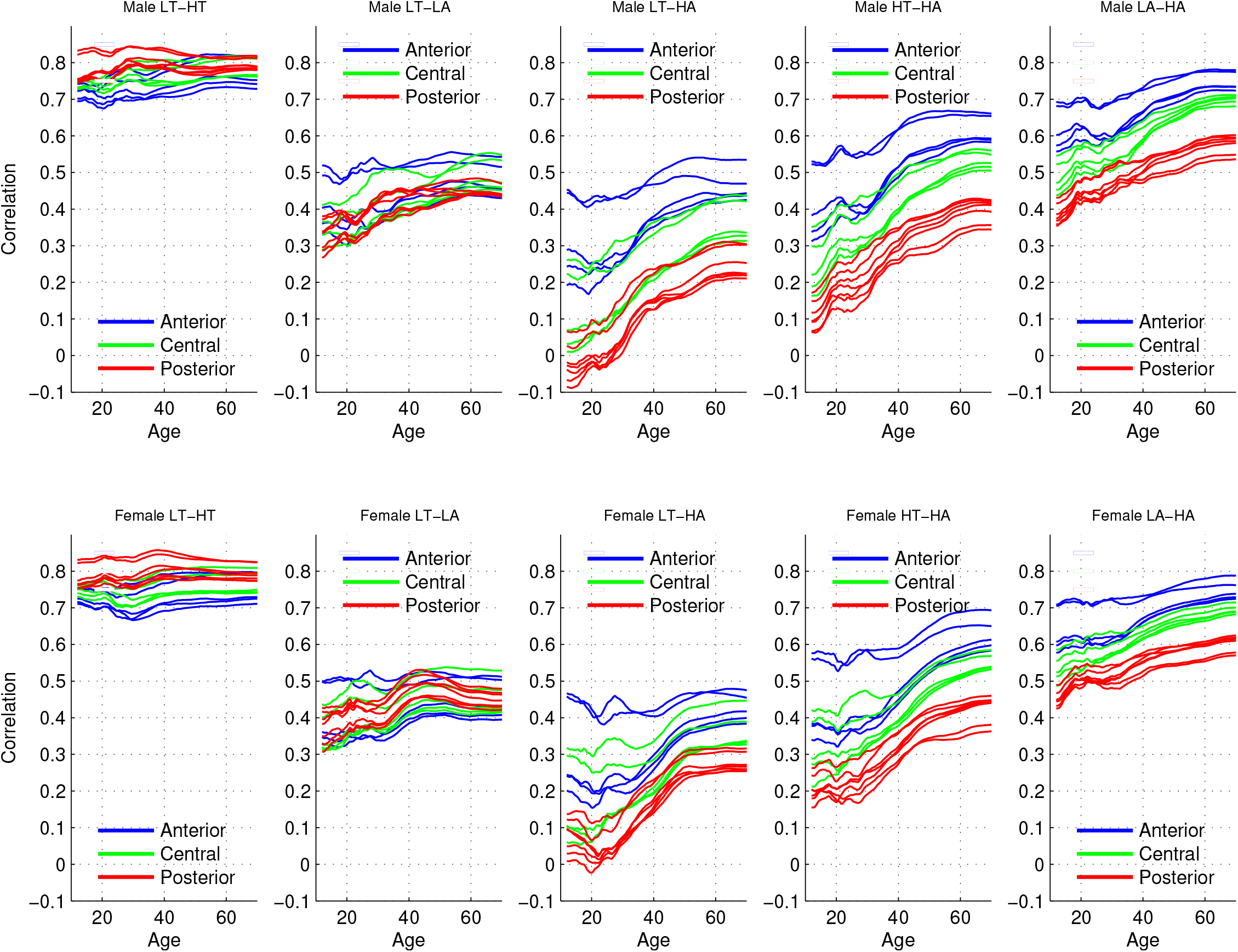
Trajectories of the male and female theta-alpha band inter-frequency correlations. Electrodes are color coded by region. Correlations are arranged symmetrically with respect to low theta and high alpha. LT = low theta … etc.

**Figure 7:**
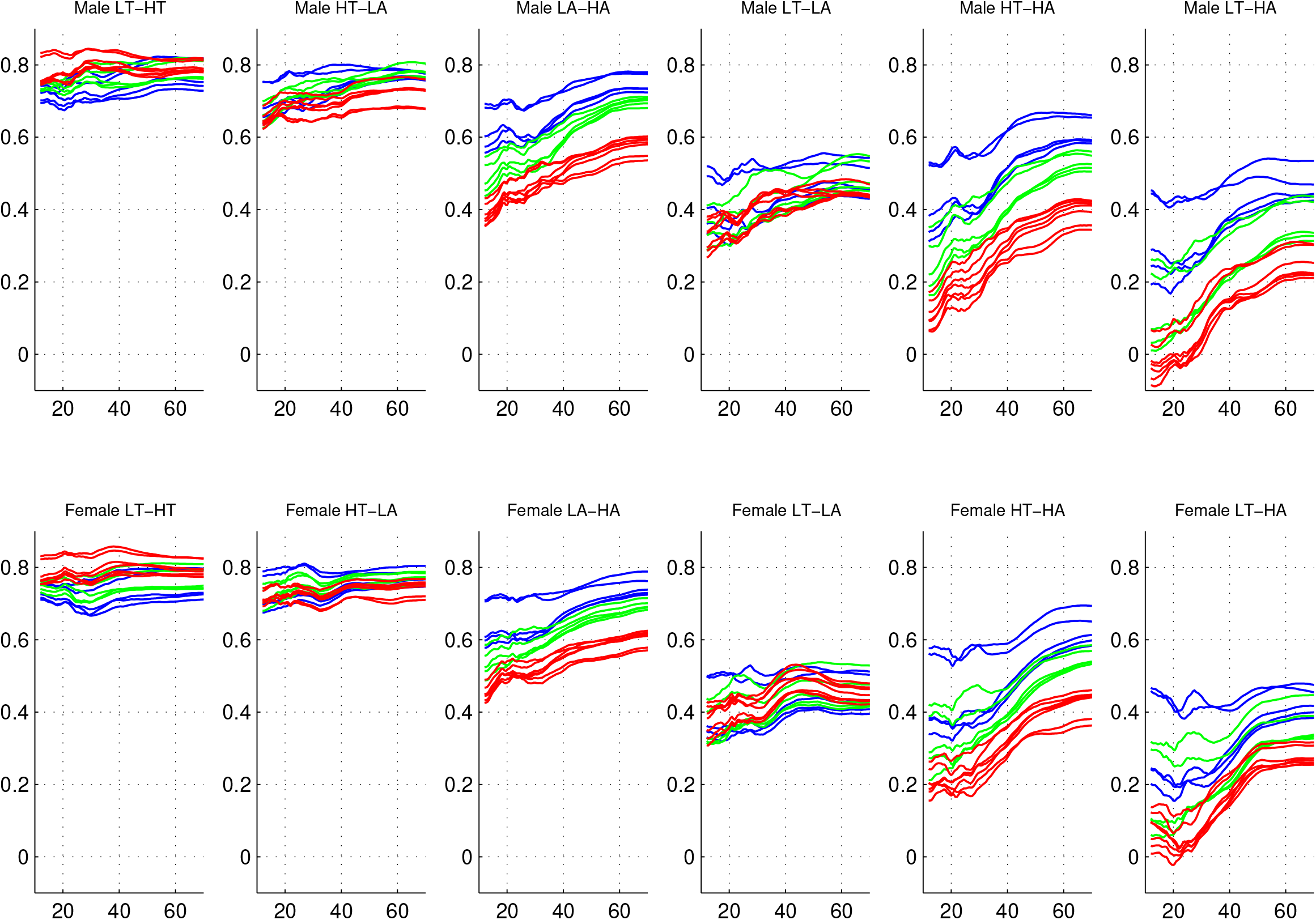
Trajectories of the male and female theta-alpha band inter-frequency correlations. Electrodes are color coded by region.

##### Alpha-Beta correlations

Inter-frequency correlations of high alpha and the three beta bands are shown in figure 8 in an order identical to the previous plot, show less between correlation difference than the theta-alpha correlations, and far less regional difference than the high alpha correlations.

**Figure 8:**
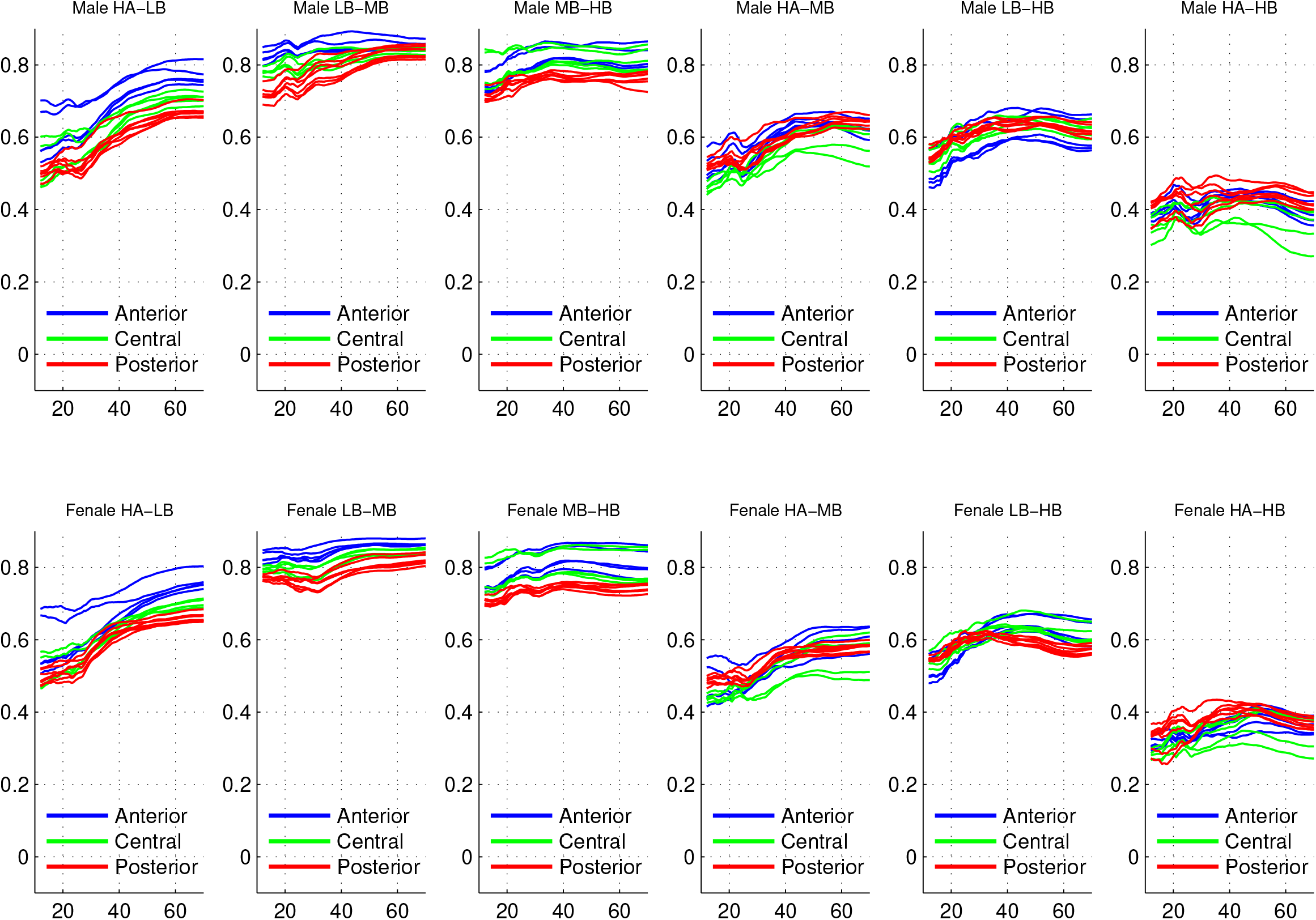
Trajectories of the male and female alpha-beta band inter-frequency correlations. Electrodes are color coded by region.

##### Alpha correlations

The pervasive increase with age of the correlation of high alpha power with the power of all the other frequency bands except for high beta is displayed in figures 9 and 10. Since this is just another way of displaying the data shown in the previous figures, the same characteristics are present, notably the regional differentiation in the theta correlations which is absent in the beta correlations. The slightly elevated levels for adjacent rows are somewhat higher vales for the peripheral electrodes.

**Figure 9:**
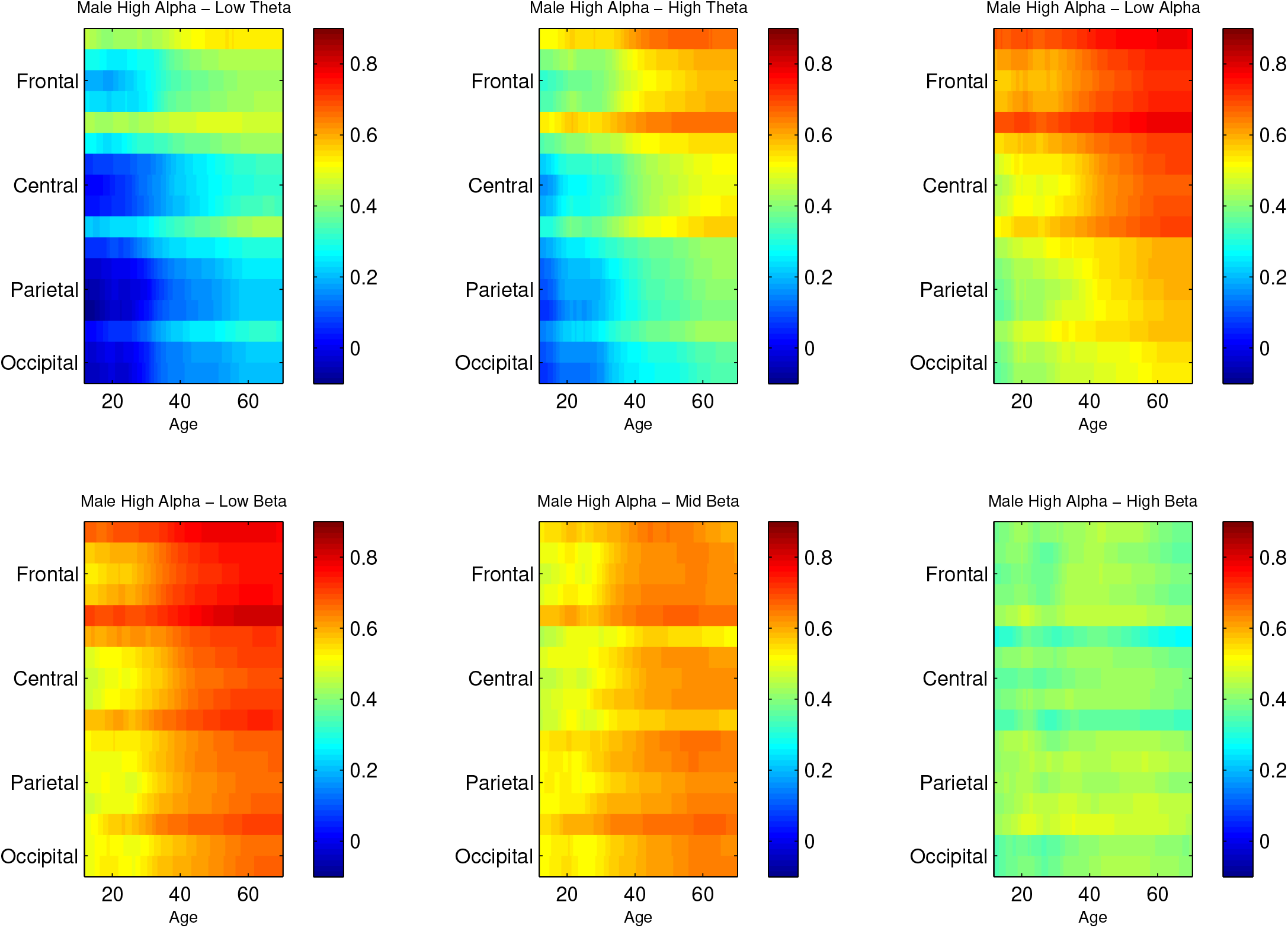
Age increase of High Alpha correlations in males. The 17 elsectrodes are arranged in order on the y-axis, age on the x-axis, and the correlation values are color-coded.

**Figure 10:**
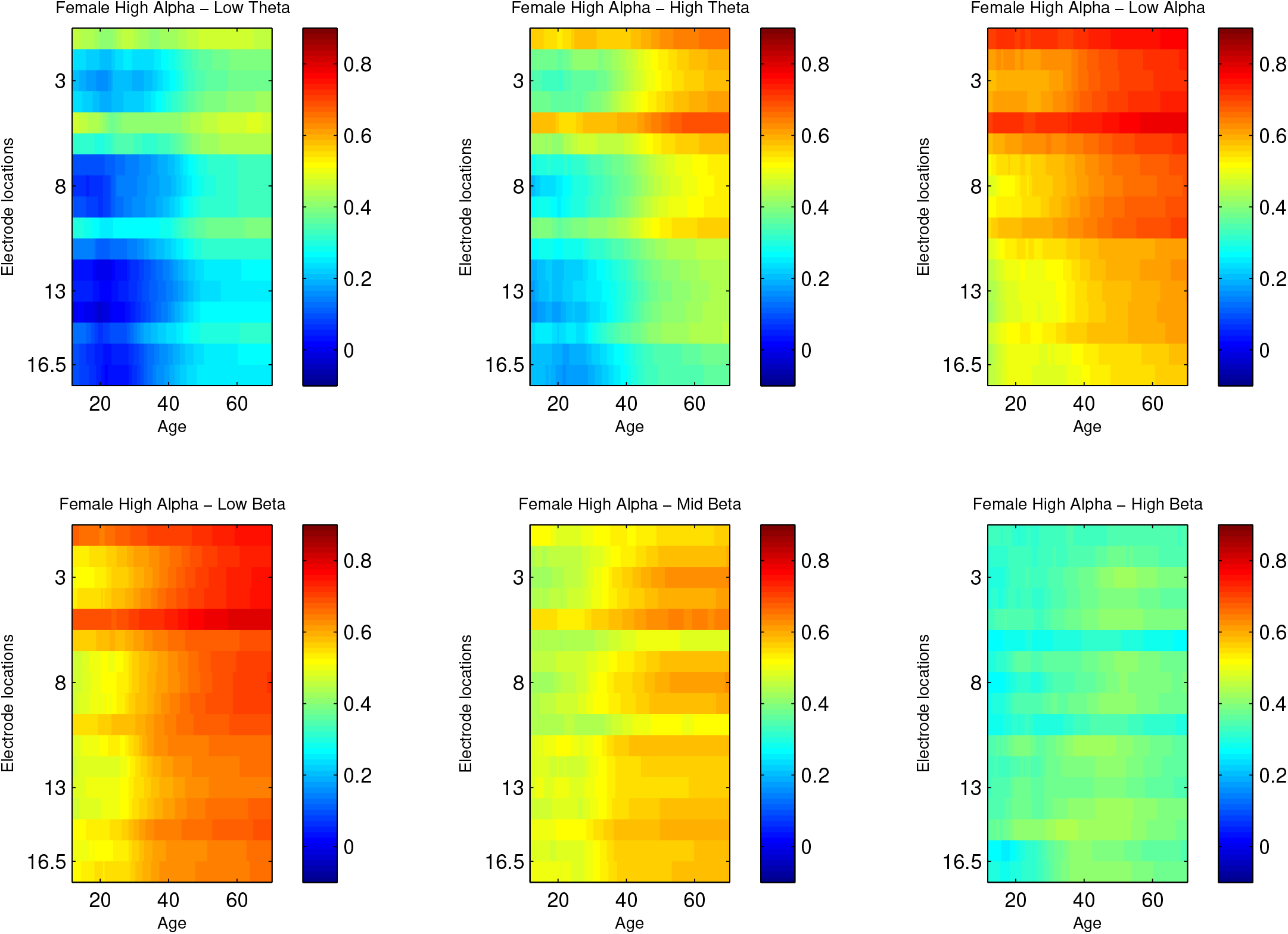
Age increase of High Alpha correlations in females. The 17 elsectrodes are arranged in order on the y-axis, age on the x-axis, and the correlation values are color-coded.

**Figure 11:**
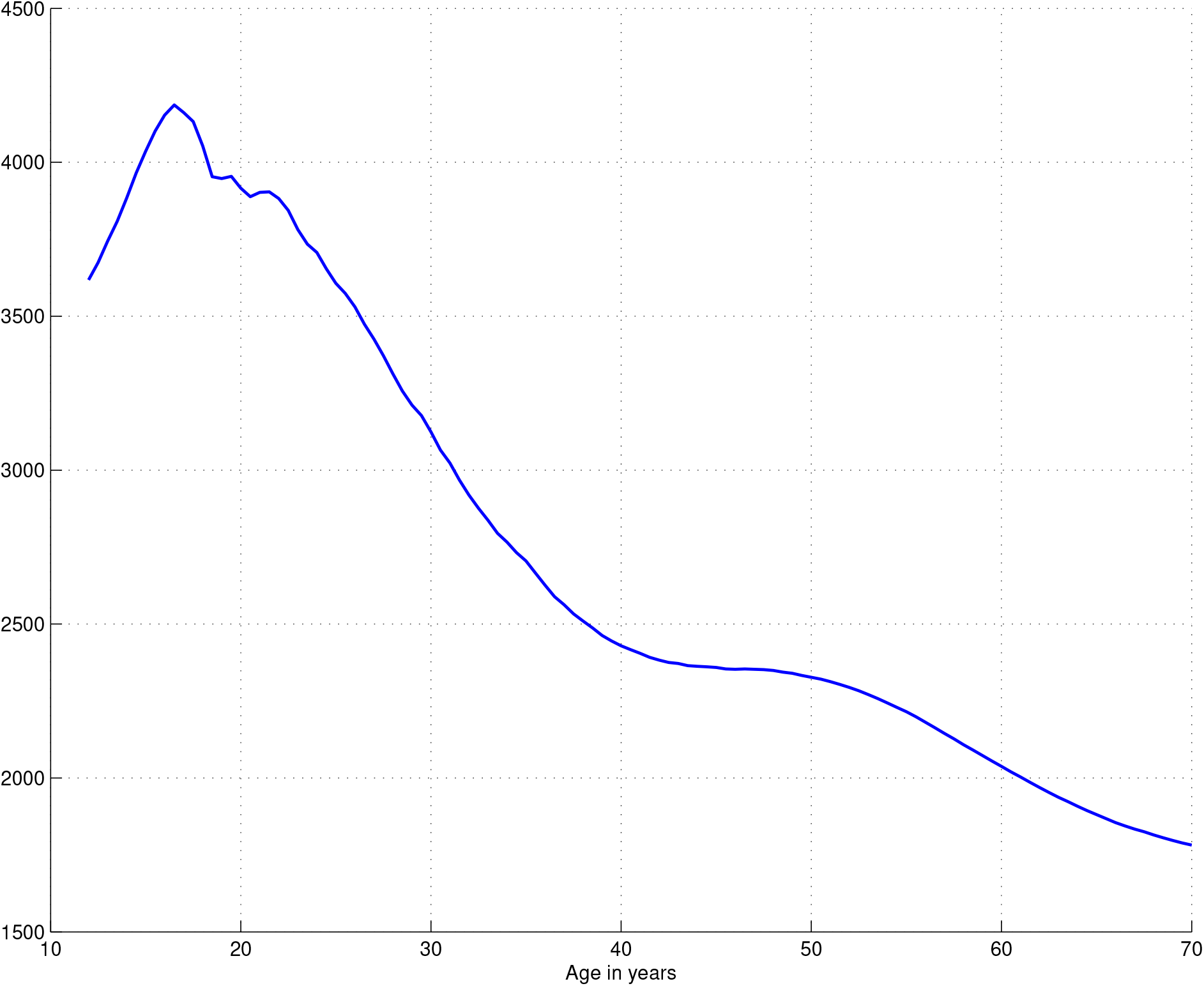
Effective sample size for bandwidth = .6

#### 3.2.2 Intra-Frequency correlation

Only the 15 inter-regional pairs and the 30 intra-regional pairs of the 136 intra-frequency correlations per frequency band were examined in order to determine whether they displayed frequency or regional characteristics comparable to those found in the inter-frequency correlation. Such characteristics were not found. Visual inspection revealed trajectories of generally constant value over age, with values consistent with the spatial relations of the electrodes in the pairs. The patterns were similar across frequencies. To check the visual impression of constancy, the coefficient of variation across the correlation values of the each of the trajectories for the four theta and alpha frequency bands was calculated. The maximum coefficient of variation over all pairs and all frequencies was less than .07, and when the maxima within the 6 regional groups in the four frequency bands were examined most values were less than .03.

### 3.3 Extensions of this analysis

The next stage of analysis is envisioned as the direct analysis of the covariance matrices, based on the methods in Chorlian et al. (2024), to provide trajectories of larger aggregates, such as encompassing all frequency bands for individual electrodes or all electrodes for individual frequency bands. As a preliminary test, the multi-dimensional scaling method of trajectory modeling from the above mentioned paper was applied to the individual trajectories of the inter-frequency covariance matrices of each of the 17 electrodes. The results determined that their first components, which accounted from 70% to over 90% of the variance for each electrode, were linear, that is, the distances between successive ages were about the same for all ages and the distance between two ages was proportional to the difference in the ages. This means that regional differences of inter-frequency covariance are variations of similar developmental processes, not variations between developmental processes with different regional/age characteristics.

The presence of the increased correlation with age of high alpha values with other frequency band values suggest that the study of the interaction of alpha production with other frequencies on an individual basis, as suggested in Becker et al. (2018), would be an informative extension of this study.

## 4 Electrode and Frequency Structure

### 4.1 Frequency bands

Frequency bands:

1. Low theta 3.0-5.0 Hz. 2. High theta 5.0-7.0 Hz. 3. Low alpha 7.0-9.0 Hz. 4. High alpha 9.0-12.0 Hz. 5. Low beta 12.0-16.0 Hz. 6. High beta 16.0-20.0 Hz. 7. High beta 20.0-28.0 Hz.

### 4.2 Regional structure of Electrodes

Frontal:

1. F7; 2. F3; 3. FZ; 4. F4; 5. F8;

Central:

6. T7; 7. C3; 8. CZ; 9. C4; 10. T8;

Parietal:

11. P7; 12. P3; 13. PZ; 14. P4; 15. P8;

5

Occipital:

16. O1; 17. O2;

## 5 Figures

The initial comprehensive views provide the trajectories of the power values across the entire age range, the entire set of frequency bands, and the entire set of electrodes for males and female in separate plots. These are followed by the derivatives of those trajectories with males and females in the same plots, beginning with the entire age range and two subsequent age ranges to better understand the variation of the onset of the male/female divergence in different frequency bands. Following these are inter-frequency correlation plots, with frequency bands labeled by pairs of letters, i.e, “LT” = low theta etc. Male and female values are on the same plots. Then follow a set of image plots, in which values are color coded, as opposed to the positional coding in the trajectory plots, to illustrate specific age, electrode and frequency band relations.

## 6 Acknowledgments

The Collaborative Study on the Genetics of Alcoholism (COGA), Principal Investigators B. Porjesz, V. Hesselbrock, A. Agrawal; Scientific Director, A. Agrawal; Translational Director, D. Dick, includes ten different centers: University of Connecticut (V. Hesselbrock); Indiana University (H.J. Edenberg, T. Foroud, Y. Liu, M.H. Plawecki); University of Iowa Carver College of Medicine (S. Kuperman, A. Anderson); SUNY Downstate Health Sciences University (B. Porjesz, J. Meyers); Washington University in St. Louis (L. Bierut, A. Agrawal, S. Hartz); University of California at San Diego (M. Schuckit); Rutgers University (D. Dick, R. Hart, J. Salvatore, J. Tischfield); The Children’s Hospital of Philadelphia, University of Pennsylvania (L. Almasy); Icahn School of Medicine at Mount Sinai (A. Goate, P. Slesinger); and Howard University (D. Scott). Other COGA collaborators include: C. Holzhauer, M. Hesselbrock (University of Connecticut); D. Lai, J. Nurnberger Jr., L. Wetherill, X., Xuei, S. O’Connor, (Indiana University); J. Kramer (University of Iowa), G. Chan (University of Iowa; University of Connecticut); C. Kamarajan, A. Pandey, D.B. Chorlian, P. Barr, S. Kinreich, G. Pandey, Z. Neale, S., C. Chatzinakos, J. Zhang, Saenz deViteri, R. Christian, A. Bingly (SUNY Downstate); G. Pathak (Icahn School of Medicine at Mount Sinai); A. Anokhin, K. Bucholz, F. Dong, A. Hatoum, E. Johnson, V. McCutcheon, J. Rice, S. Saccone (Washington University); F. Aliev, Z. Pang, S. Kuo, S. Brislin, J. Moore (Rutgers University); A. Merikangas (The Children’s Hospital of Philadelphia and University of Pennsylvania); M. Gitik, NIAAA Staff Collaborator. We continue to be inspired by our memories of Henri Begleiter and Theodore Reich, founding PI and Co-PI of COGA, and also owe a debt of gratitude to other past organizers of COGA, including Ting-Kai Li, P. Michael Conneally, Raymond Crowe, and Wendy Reich, for their critical contributions. This national collaborative study is supported by NIH Grant U10AA008401 from the National Institute on Alcohol Abuse and Alcoholism (NIAAA) and the National Institute on Drug Abuse (NIDA).

